# *Cis*- and *trans*-acting factors influence expression of the *norM*-encoded efflux pump of *Neisseria gonorrhoeae* and levels of gonococcal susceptibility to substrate antimicrobials

**DOI:** 10.1101/308031

**Authors:** Corinne E. Rouquette-Loughlin, Vijaya Dhulipala, Jennifer L. Reimche, Erica Raterman, Afrin A. Begum, Ann E. Jerse, William M. Shafer

**Author notes:** Address correspondence to WM Shafer.

## Abstract

The gonococcal NorM efflux pump exports substrates with a cationic moiety including quaternary ammonium compounds such as berberine (BE) and ethidium bromide (EB) as well as antibiotics such as ciprofloxacin and solithromycin. The *norM* gene is part of a four gene operon that is transcribed from a promoter containing a polynucleotide tract of 6 or 7 thymidines (Ts) between the -10 and -35 hexamers; the majority of gonococcal strains analyzed herein contained a T-6 sequence. Primer extension analysis showed that regardless of the length of the poly-T tract, the same transcriptional start site (TSS) was used for expression of *norM*. Interestingly, the T-6 tract correlated with a higher level of both *norM* expression and gonococcal resistance to NorM substrates BE and EB. Analysis of expression of genes downstream of *norM* showed that the product of the *tetR*-like gene has the capacity to activate expression of *norM* as well as *murB*, which encodes an acetylenolpyroylglucosamine reductase predicted to be involved in the early steps of peptidoglycan synthesis. Moreover, loss of the TetR-like transcriptional regulator modestly increased gonococcal susceptibility to NorM substrates EB and BE. We conclude that both *cis*- and *trans*-acting regulatory systems can regulate expression of the *norM* operon and influence levels of gonococcal susceptibility to antimicrobials exported by NorM.

## INTRODUCTION

*Neisseria gonorrhoeae* is a strict human pathogen and is the etiologic agent of the sexually transmitted infection (STI) termed gonorrhea, which is the second most prevalent bacterial STI in the United States and had a world-wide incidence in 2012 of an estimated 78 million infections (1). The gonococcus has adapted numerous strategies to survive attack by antimicrobials including classical antibiotics used in treatment of infection and those of host origin that participate in innate host defense. In this respect, gonococci use efflux pumps to resist the antimicrobial action of beta-lactam and macrolide antibiotics as well as cationic antimicrobial peptides and long-chain fatty acids (2–4). The capacity of gonococci to utilize efflux pumps to resist clinically useful antibiotics is of interest given the emergence of strains resistant to current and past front-line antibiotics (2, 5–7). The contribution of efflux pumps in aiding bacterial evasion of antimicrobials can be enhanced by mutations that de-repress expression of efflux pump-encoding genes (8). With respect to gonococci, previous work revealed that overexpression of the *mtrCDE* efflux pump operon due to *cis*- or *trans*-acting mutations can contribute to clinically relevant levels of antibiotic resistance (9, 10) and increase bacterial fitness during experimental infection of the lower genital tract of female mice presumably due to enhanced resistance to host antimicrobials (11, 12).

In this study, we investigated the regulation of the gonococcal *norM* gene. NorM belongs to the Multidrug And Toxic compound Extrusion (MATE) family of efflux proteins which are Na^+^- or H^+^-coupled transporters and are universally present in all living organisms (13). Gonococcal NorM is highly similar (56%) to NorM of *Vibrio parahaemolyticus* (14). We previously reported that NorM can export substrates with a cationic moiety including berberine (BE), ciprofloxacin (CIP) and ethidium bromide (EB) (15). Additionally, loss of the NorM efflux pump in multi-drug resistant strain H041 was found by Golparian *et al*. (6) to increase gonococcal susceptibility to solithromycin. Herein, we investigated *cis*- and *trans*-acting regulatory mechanisms that influence *norM* expression and the consequence of such on antimicrobial resistance. Importantly, we identified a heretofore undescribed TetR-like regulator that activated the *norM* gene as well as a single-base-pair deletion that resulted in a stronger *norM* promoter.

## RESULTS and DISCUSSION

### *Cis*-acting transcriptional regulation of *norM* in *N. gonorrhoeae* and influence on antimicrobial resistance

Bioinformatic analysis (http://www.ncbi.nlm.nih.gov) indicated that *norM* (NGO0395) is the first gene of an operon that also contains three downstream genes annotated as *murB* (NGO0394), which encodes a putative UDP-N-acetylenolpyruvoylglucosamine reductase involved in the initial steps of the peptidoglycan synthesis (Fig. 1A), NGO0393, which encodes a TetR-like family transcriptional regulator homolog, and NGO0392 which encodes a hypothetical protein. (Fig. 1A). Using total RNA prepared from strain FA19Str^R^ in RT-PCR experiments, we confirmed transcriptional linkage of *norM* and *murB* as well as *murB* and *tetR* (data not shown), which supports the hypothesis that the genes form an operon. Primer extension analysis of this RNA indicated the presence of 2 distinct TSSs. One TSS was located upstream of *norM* that corresponded to that described previously by Rouquette-Loughlin *et al*. (15) as well as another TSS located upstream of *tetR*. This result suggests the presence of two distinct promoters that express genes within the operon with one capable of directing transcription of the entire operon and a second driving the transcription of *tetR* and possibly NGO0392 (Fig. 1B).

**FIG 1.**
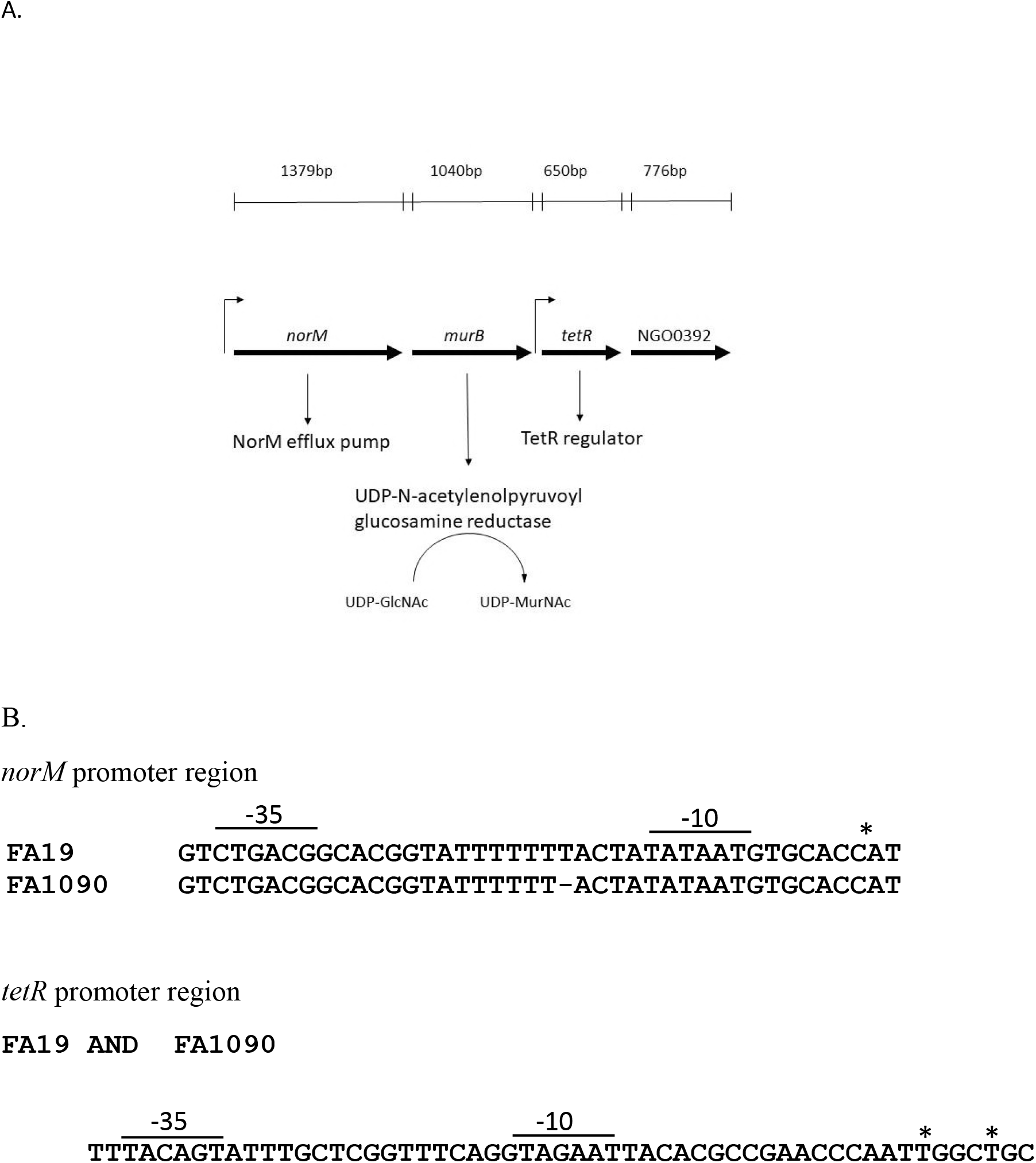
A. The organization of the *norM* operon is depicted. The length and transcriptional direction (arrows) of the genes are shown. B. Sequences of the *norM* and *tetR* promoter regions from strains FA19 and FA 1090. The -10 and -35 hexamers are indicated, and * represents the TSS.

DNA sequencing of the *norM* promoter region of strain FA19Str^R^ revealed the presence of a stretch of 7 Ts between the -10 and -35 hexamers (Fig.1B). In order to learn if this poly-T stretch is common amongst gonococci, we performed a bioinformatic analysis of a 200 bp region upstream of the *norM* translational start codon using thirty-one gonococcal whole genome sequences that are available on-line (http://www.ncbi.nlm.nih.gov). This analysis revealed that the majority (77.4 %) of gonococcal strains had a stretch of 6 Ts (including strains FA1090 and certain WHO reference strains) while a minority (22.6%) of strains had 7 Ts (including strains FA19 and MS11). Using a PCR-generated product, we also sequenced this *norM* upstream region from ten clinical isolates and found that nine of ten had the T-6 sequence (Table S1). Thus, we conclude that the T-6 sequence predominates in gonococci. In contrast, our analysis of whole genome sequences of eighty-six *N. meningitidis* strains that are publicly available (http://www.ncbi.nlm.nih.gov) revealed that eight-five (99%) have a *norM* promoter with a T-7 repeat sequence (data not presented).

Despite the difference in T repeat length, primer extension analysis revealed the same TSS positioned upstream of the *norM* promoter was possessed by strains FA19 (T-7) and FA 1090 (T-6), which was identified as a C residue located 6 bp downstream of the -10 hexamer (data not presented; summarized in Fig. 1B). The level of the *norM* transcript in strains FA19Str^R^ and FA1090 was determined by quantitative qRT-PCR (qRT-PCR) analysis using total RNA prepared from mid- and late-logarithmic cultures, which showed that the *norM* transcript was 2.4-and 4.2-fold higher in strain FA1090 compared to that of FA19Str^R^ at mid- and late-logarithmic phases, respectively (Fig. 2). Previous studies on the regulation of the *mtrCDE* efflux pump-encoding operon revealed that the distance between the -10 and -35 promoter hexamers can significantly influence transcription and levels of gonococcal resistance to antimicrobials exported by MtrCDE (9, 16). Guided by this work, we constructed *norM* mutants of FA1090 and FA19Str^R^ by insertional inactivation with the non-polar *aphA-3* cassette and found that while loss of *norM* influenced gonococcal susceptibility to NorM substrates (BE and EB), the impact was greatest in strain FA1090 (T-6 promoter) compared to strain FA19Str^R^ (T-7 promoter) (Table 1). In order to determine if inactivation of *norM* would increase susceptibility of a more recent gonococcal clinical isolate displaying resistance to multiple antibiotics (6), we constructed a *norM::kan* transformant of strain H041 (T-6 promoter-Table S1). We found that loss of the NorM efflux pump decreased resistance of H041 to both BE and EB (Table 1) as well as solithromycin (4-fold decrease in MIC; data not presented).

**FIG 2.**
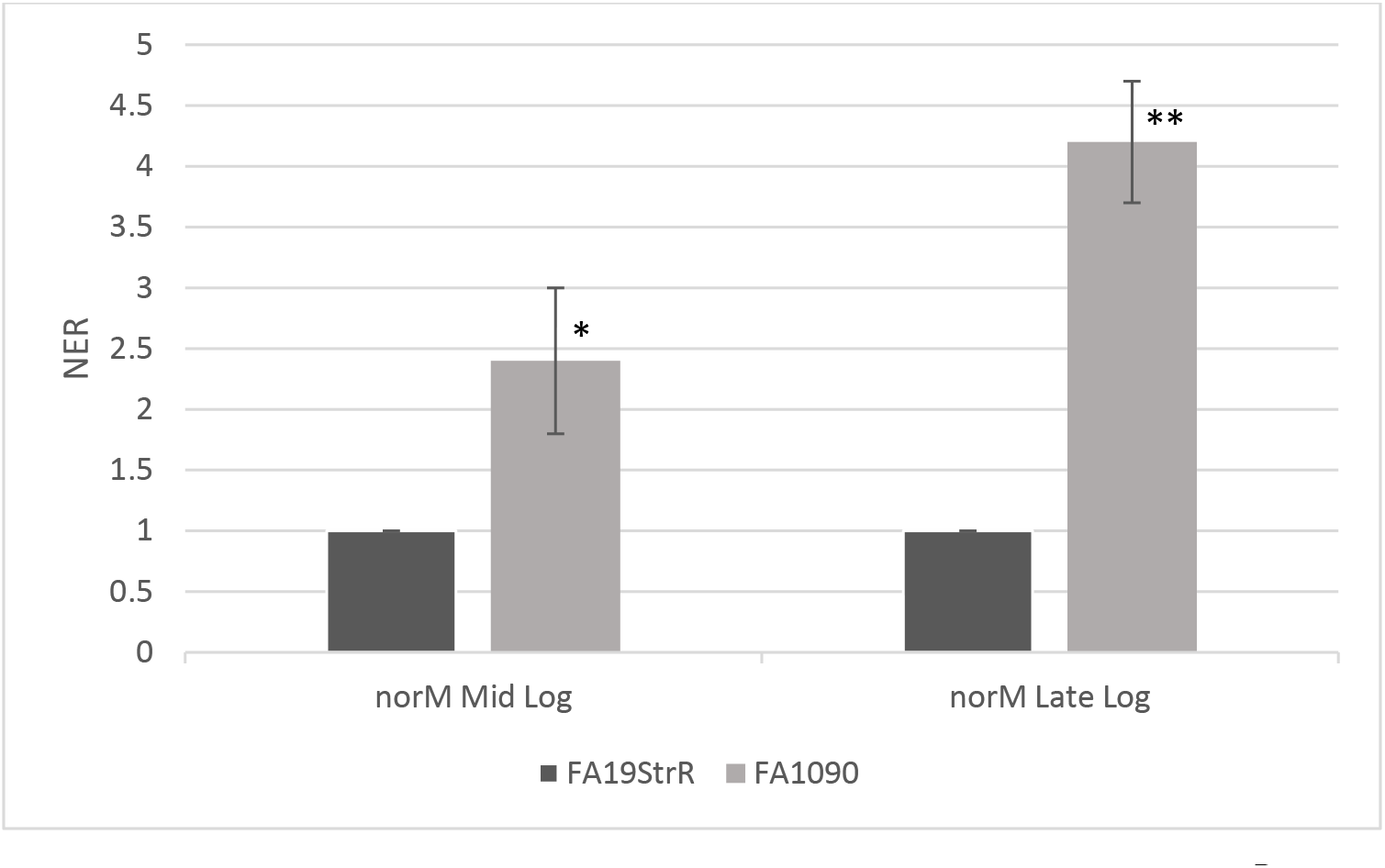
Quantitative RT-PCR results with *norM* in strains FA19Str^R^ and strain FA 1090 at mid- and late-logarithmic phases of growth. Error bars represent standard deviations from the means of 3 independent experiments. Normalized Expression Ratios (NER) were calculated using 16S rRNA expression. \**p*=0.018, \*\**p*=0.004 for comparison of values of FA1090 vs. FA19Str^R^. The statistical significance was determined by Student’s *t*.-test.

**TABLE 1.** Susceptibility of gonococcal strains to NorM substrates

| MIC (µg/ml)<sup>a</sup> |  |  |
| --- | --- | --- |
| Strains | BE<sup>b</sup> | EB |
| FA19Str<sup>R</sup> | 5 | 1.25 |
| FA19Str<sup>R</sup>*norM::kan* | 1.25 | 0.625 |
| FA19Str<sup>R</sup>*norM::kanC3* | 5 | 1.25 |
| FA19Str<sup>R</sup>*tetR::kan* | 2.5 | 0.625 |
| FA19Str<sup>R</sup>*tetR::kanC4* | 5 | 1.25 |
| FA19Str<sup>R</sup><sub>P<sub>norM</sub>FA1090</sub> | 10 | 2.5 |
| FA 1090 | 20 | 5 |
| FA 1090*norM::kan* | 1.25 | 0.625 |
| FA 1090*tetR::kan* | 20 | 5 |
| HO41 | >40 | 20 |
| HO41*norM::kan* | 10 | 1.25 |

<sup>a</sup>All results are representative from 3 or more independent determinations
<sup>b</sup>BE: berberine; EB: Ethidium bromide

In order to determine the influence of the *norM* promoter T repeat sequence on gonococcal expression of the *norM* operon and resistance to NorM substrates, we exchanged the *norM* promoter region of FA19Str^R^ with that of FA1090 by transformation. DNA sequencing of a PCR fragment from a representative transformant strain (FA19Str^R^ _PnorMF1090_) confirmed the presence of the T-6 instead of T-7 repeat element (data not presented). Importantly, FA19Str^R^ _PnorMF1090_ displayed a 5-fold increase in expression of *norM* as assessed by qRT-PCR (Fig. 3) and displayed a two-fold increase in resistance to EB and BE compared to parental strain FA19Str^R^ (Table 1). These combined results indicated that the length of the T-track can influence levels of gonococcal expression of *norM* and resistance to NorM substrates.

**FIG 3.**
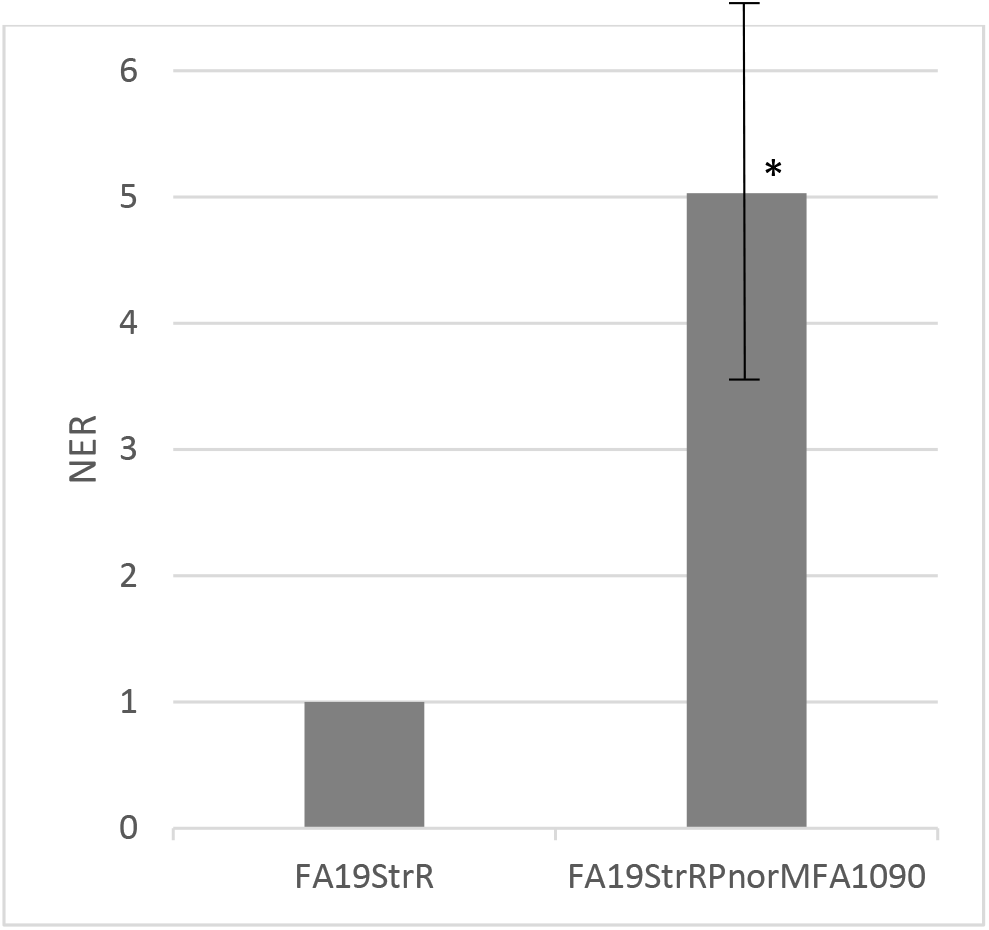
Quantitative RT-PCR results with *norM* in strains FA19Str^R^ and strain FA19Str^R^_PnorM1090_ at mid -logarithmic phase of growth. Error bars represent standard deviations from the means of 3 independent experiments. Normalized Expression Ratios (NER) were calculated using 16S rRNA expression. *p=0.01. The statistical significance of the results was determined by Student’s *t*.-test.

### *Trans*-acting transcriptional regulation of *norM* and influence on antimicrobial resistance

Bioinformatic analysis revealed that the putative TetR-like protein (216 amino acids) encoded by a gene within the *norM* operon is highly conserved in gonococci. This finding is exemplified by 100% amino acid identity of the protein that would be produced by strains FA19 and FA1090; the protein is also highly similar (97% identical) to a counterpart encoded by meningococci (data not presented). That this TetR-like protein can act as a transcriptional regulator was suggested by the presence of a helix-turn-helix DNA-binding domain at the N-terminus (amino acids 15-61). Further, the position of *tetR* downstream of *norM* suggested that the TetR-like protein might exert transcriptional control of *norM* and other genes (e.g., *murB*) in the operon. In order to test this possibility, *tetR::kan* mutants of strains FA19Str^R^ and FA1090 were constructed and analyzed for changes in susceptibility to NorM substrates and levels of gene transcripts within the operon. We noted that with strain FA19Str^R^, but not FA1090, insertional inactivation of *tetR* reproducibly resulted in two-fold decrease in gonococcal resistance to BE and EB (Table 1). Although the impact of loss of the TetR-like protein was modest, complementation of the FA19Str^R^ *tetR::kan* strain with a pGCC4 construct bearing the wild-type *tetR* gene expressed at the *aspC-lctP* locus from an IPTG inducible *lac* promoter restored wild-type levels of antimicrobial susceptibility (Table 1).

Based on the above-described results, we used strain FA19Str^R^ to ascertain if the TetR-like protein could regulate the *norM* operon. Results from qRT-PCR experiments indicated that loss of the TetR-like protein decreased expression of both *norM* and *murB* (Fig. 4), which is consistent with the transcriptional linkage of these genes by a promoter upstream of *norM*. To investigate if the TetR homolog could directly activate transcription of the *norM* operon, a recombinant His-tagged TetR protein was purified and employed in competitive EMSA experiments that used a radiolabeled PCR probe containing 344 bp of DNA upstream of *norM*. The results from DNA-binding experiments showed that TetR could bind to the probe in a specific manner as binding could be inhibited by the unlabeled *norM* PCR product, but not by a non-specific PCR probe (Figure 5). Thus, this gene regulator serves as a transcriptional activator of the *norM* operon.

**FIG 4.**
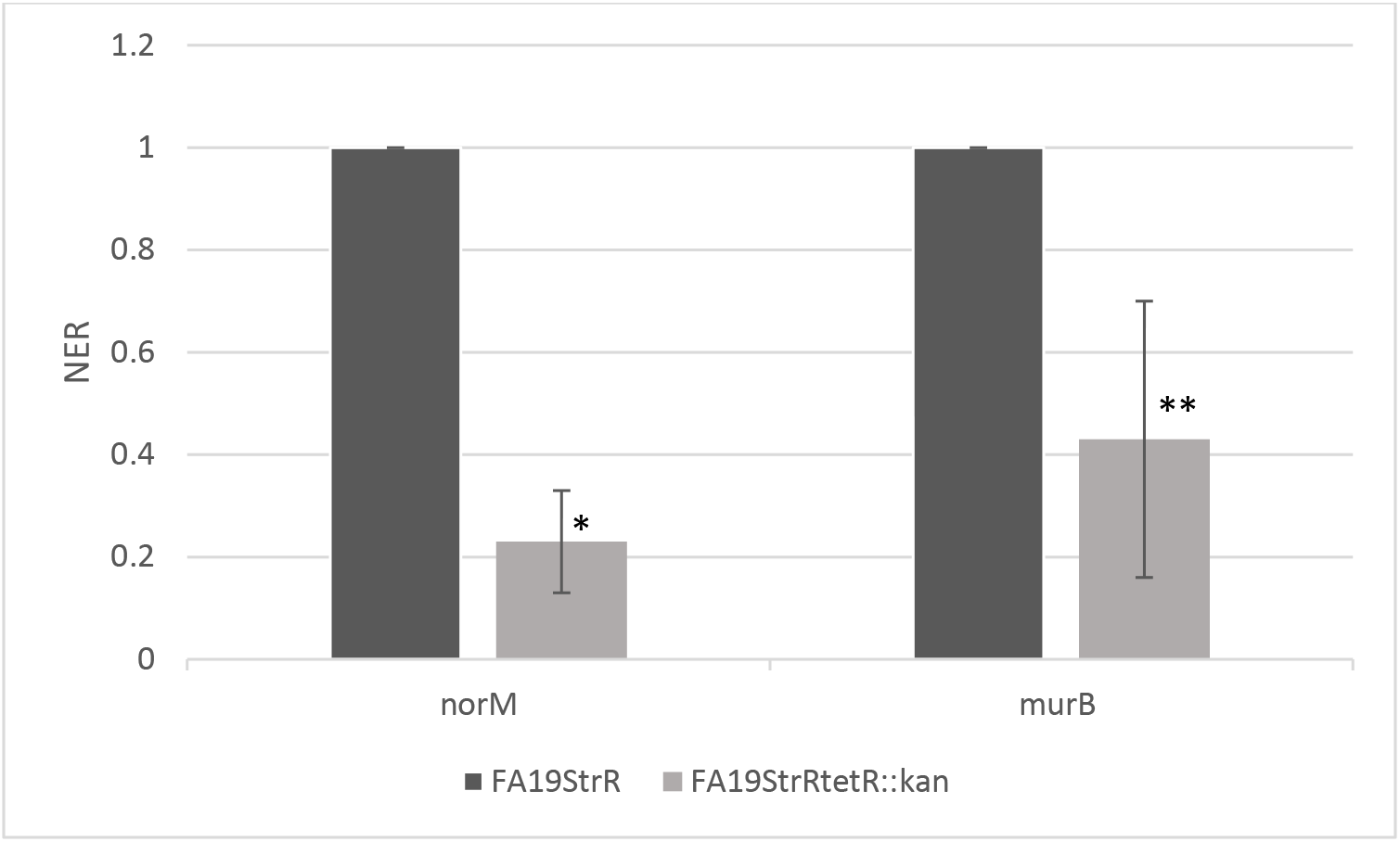
Quantitative RT-PCR results with *norM* and *murB* in strains FA19Str^R^ and strain FA19Str^R^*tetR::kan* at mid-logarithmic phase of growth. Error bars represent standard deviations from the means of 3 independent experiments. Normalized Expression Ratios (NER) were calculated using 16S rRNA expression. \**p*=0.0004, \*\**p*=0.022 for comparison of values of FA19Str^R^*tetR::kan* vs. FA19Str^R^. The statistical significance of the results was determined by Student’s *t*.-test.

**FIG 5.**
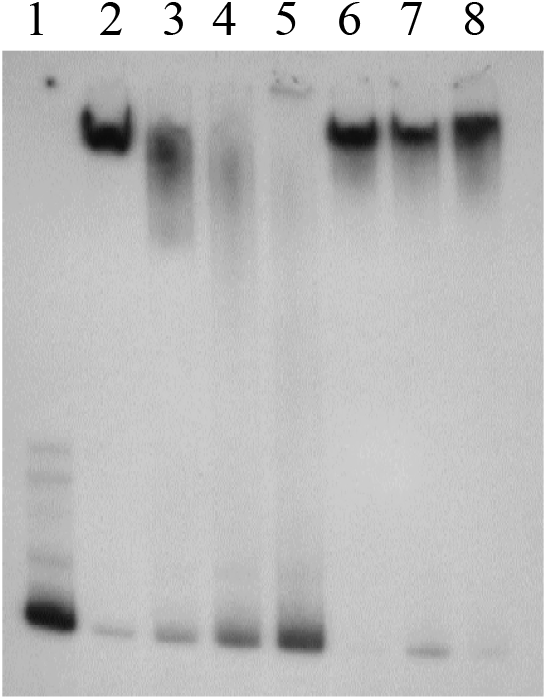
Competitive EMSA demonstrating binding specificity. The purified TetR-His protein binds to the *norM* promoter from strain FA19 in a specific manner. Lane 1, hot probe N11/N14* alone; lane 2, hot probe N11/N14* plus 8 μg of TetR-His; lanes 3 to 5, hot probe N11/N14* plus 8 μg of TetR-His plus 100X, 200X and 400X respectively of unlabeled N11/N14; lanes 6 to 8, hot probe N11/N14* plus 8 μg of TetR-His plus 100X, 200X and 400X respectively of unlabeled *rnpB*.

As a member of the MATE family of efflux pumps, the gonococcal NorM efflux pump has the capacity to export antimicrobial quartenary ammonium compounds (ref. 15 and Table 1). The conservation of *norM* among gonococci suggests a role for NorM in the survival of gonococci. Thus, we hypothesized that NorM might also export host-derived antimicrobials and promote survival of gonococci during infection. However, using the established female mouse model of lower genital tract infection previously employed to determine the biological significance of the gonococcal MtrCDE efflux pump and cognate regulatory systems (11, 17), we did not detect a fitness or survival defect of gonococci (FA19Str^R^ and FA1090) bearing a null mutation in *norM* when competed with the wild-type parent strains (Figure S1). It is important to note that this model may not fully recapitulate the repertoire of antimicrobials present at human female or male mucosal surfaces. Moreover, the infection model employed is limited to the lower genital tract of female mice and distinct antimicrobials in the upper tract that might serve as NorM substrates could exist. For instance, differences in the presence and level of antimicrobial peptides have been reported at mucosal surfaces of the human ectocervix and endocervix (18). Hence, the possibility that NorM promotes survival of gonococci during human infection by promoting resistance to a host antimicrobial cannot be discounted.

As with other bacterial efflux pump-encoding genes (8, 9, 16), we conclude that the gonococcal *norM* gene is subject to transcriptional control that influences its expression and levels of bacterial resistance to antimicrobials that can be exported by NorM. It is of interest that both *cis*- and *trans*-acting regulatory processes identified in this study can modulate expression of *norM* and that these regulatory schemes seem dependent on the length a poly-T tract in the *norM* promoter. The majority of gonococci contain a T-6 tract in the promoter that seems to enhance transcription of *norM*. In contrast, strain FA19, which we have used extensively in our work on gonococcal efflux pumps (3, 4, 15–19), is representative of the minority of gonococcal strains harboring a T-7 sequence. Since mutations that increase or decrease spacing between the -10 and -35 hexamers can influence the fidelity of gene expression due to impacting interactions of RNA polymerase, as has been observed with nucleotide deletions or insertions within the *mtrR* promoter (16, 19), it is likely that the single T difference can impact *norM* expression in gonococcal strains with a T-6 or T-7 sequence by influencing promoter recognition by RNA polymerase.

In addition to this *cis*-acting regulatory mechanism, the TetR-like protein encoded by a gene within the *norM* operon can influence expression of *norM* in strain FA19. Importantly, the TetR DNA-binding protein also activates expression of *murB*, which is consistent with its transcriptional linkage with *norM*. It is of interest that a gene (*murB*) encoding an enzyme involved in the earliest steps of peptidoglycan biosynthesis (19) is co-regulated with *norM* by both *cis*- and *trans*-acting regulatory schemes. Thus, the fidelity of early stages of peptidoglycan biosynthesis may be modulated by transcriptional control systems that also influence expression of *norM* and levels of gonococcal resistance to antimicrobials exported by NorM.

The chemical characteristics of known substrates of the gonococcal NorM efflux pump suggest that the clinical efficacy of future antimicrobials having similar properties may be influenced by constitutive or inducible changes in *norM* expression. We hypothesize that increased expression of *norM* coupled with other mutations could result in clinical resistance to antibiotics used in the future for treatment of gonorrhea. Based on earlier work with multidrug resistant strain H041 by Golparian et al (6) and our findings with this clinical isolate, this possibility should be considered for solithromycin and its future use in treatment of gonorrhea. In a broader sense, de-repression of bacterial efflux pump genes due to constitutive mutations as well as inducible activation systems should be considered as a contributing factor by which gonococci (or other bacteria) might develop clinical resistance to antibacterials under development.

## MATERIALS AND METHODS

### Gonococcal strains, growth conditions, and determination of susceptibility to antimicrobial agents

Strains FA19, FA19Str^R^ and FA 1090 were the primary gonococcal strains used in this study. These strains and their genetic derivatives as well as their susceptibility to antimicrobials are presented in Table 1. We also sequenced the *norM* promoter region from ten clinical isolates (Table S1, see below). Gonococcal strains were grown overnight at 37°C under 5 % (v/v) CO_2_ on GCB agar containing defined supplements I and II (9). Susceptibility of test strains to antibiotics was performed by the agar dilution method and reported as the minimal inhibitory concentration (MIC) (21). IPTG was added a final concentration of 1 mM to MIC plates to allow complementation by the pGCC4 vector (22). Antibiotics were purchased from Sigma Chemical Co. (St. Louis, MO). Solithromycin was obtained from Med Chem Express (Monmouth, NJ). *Escherchia coli* strains were grown overnight at 37°C on LB agar.

### Sequencing of the *norM* promoter

The *norM* promoter region was PCR-amplified from gonococci using primers norMPac1 (5’-GATCTTAATTAACAATGCCGTCAAGTCGTTAAA-3’) and N10 (5’-CATCACGGTATCGACGAAACGATGCCC-3’). The resulting PCR product was sequenced using norMPac1.

### Construction of the *norM* and *tetR*-negative mutants and their complemented strains

The pBAD*norM::kan* construct (15) was transformed into FA19Str^R^ and transformants were selected on GC agar supplemented with 50 μg/ml of kanamycin (Kan). FA19Str^R^ *norM::kan* transformants were verified by PCR and sequencing. The pGCC3 vector (22) was used to complement FA19Str^R^*norM::kan*. This complementation system allows the integration of a wild-type copy of *norM* under its own promoter at the transcriptionally silent intergenic region between *lctP* and *aspC*. norMPac1 and norMpme1 (5’-GATCGTTTAAACTATCGGATGGGTTGCATGGT -3’) were used to amplify the *norM* gene and its own promoter. The resulting PCR product was cloned into the pGCC3 vector. The pGCC3norM construct was verified by sequencing and then transformed into FA19Str^R^ *norM::kan*. FA19Str^R^ *norM::kan*C3 transformants were selected on GC agar plates supplemented with 1 μg/ml of erythromycin (Ery) and verified by colony PCR and sequencing. The *norM* gene from FA1090 was amplified using N6 (5’-TCGGTATCGGATGGGTTGC-3’) and N4 (5’-ATGCTGCTCGACCTCGACC-3’) primers, the resulting PCR product was cloned into pBAD. pBAD*norM* was then digested by Nae1 and a non-polar Kan-resistance cassette from pUC18K (23) was inserted. The resulting construct was transformed into FA1090 and transformants were selected on GC agar plates supplemented with 50 μg/ml of kan. FA 1090*norM::kan* transformants were verified by colony PCR and sequencing. To construct the *tetR*-negative mutant, pUC19 vector was digested by BamH1 and EcoR1 and PCR was performed on FA19 genomic DNA with E1tetR (5’-GGAATTCCTGTATGGGCAGGTTGATGTC-3’) and Sma1R (5’-TCCCCCGGGGGATCGCCCAACAATTCGGCAC-3’) primers and B1tetR (5’-CGCGGATCCGCGCTGAAGGGCTTCCAAATCGG-3’) and Sma1F (5’-TCCCCCGGGGGAACACAATACCTTTACCCAAGC-3’). The resulting PCR products were ligated into pUC9 digested with BamH1 and EcoR1 digested to create a Sma1 site 356 bp downstream the ATG of the *tetR* gene. The resulting construct was verified by PCR and then digested by Sma1. The Kan-resistance cassette was PCR-amplified with pfu using AphF (5’-GTGACTAACTAGGAGGAATAAAT-3’) and AphR (5’-GGTCATTATTCCCTCCAGGTA-3’) primers and pUC18K (22) as a template. The kan cassette was then cloned into the Sma1 digested pUC19*tetR*. The ligation reaction was transformed into *E. coli* DH5α and transformants were selected on LB agar plates supplemented with 50 μg/ml of Kan. The resulting construct was then verified by sequencing and used to transform strains FA19Str^R^ and FA1090 for resistance to Kan (50 ug/ml). The pGCC4 vector was used to complement FA19Str^R^*tetR::kan*. This complementation system allows the integration of a wild-type copy of *tetR* under an IPTG inducible promoter at the transcriptionally silent intergenic region between *lctP* and *aspC*. tetRpac1 (5’-GATCTTAATTAAAGCCTGTAAATCCAAGGAGTA-3’) and tetRpme1 (5’-GATCCGTTTAAACCGTCTGAAGGCTGATTCGG-3’) were used to amplify the *tetR* gene. The resulting PCR product was cloned into the pGCC4 vector. The pGCC4*tetR* construct was verified by sequencing and then transformed into FA19Str^R^*tetR::kan*. Transformant FA19Str^R^*tetR::kan*C4 was obtained by selection with 1 μg/ml of chloramphenicol (CMP) and the genotype was verified by colony PCR.

### Construction of FA19Str^R^ _PnorMFA1090_

Primers N5 (5’-GGATGAACATCGGCACCTTG-3’) and norMPacl (5’- GATCTTAATTAACAATGCCGTCAAGTCGTTAAA-3’) were used to amplify the *norM* promoter region of strain FA 1090. The resulting 1385 bp PCR product was then transformed into strain FA19Str^R^ and transformants were selected on GC agar containing defined supplements I and II supplemented with EB (1 μg/ml). Transformants were then verified by DNA sequencing of a PCR product generated using primers N5 and norMPacI.

### Mapping transcriptional start sites by primer extension analysis

Total RNA from strains FA19 and FA1090 was prepared at late-logarithmic phase of growth in GC broth as described above by the TRIzol method as directed by the manufacturer (Thermo Fisher Scientific, Waltham, MA). Primer extension experiments were performed as described previously (9, 16) on 6 μg of total RNA with primer N11 (5’-CGGTCAGCAGGCGGATTTCTTTCAGG-3’) for *norM* and primer tetRPE (5’-TGGCGTCGATGATGCGGG-3’) for *tetR*. The AMV Reverse Transcriptase Primer Extension transcription start sites (TSSs) were determined by electrophoresis of the extension products on a 6% (w/v) DNA sequencing acrylamide gel adjacent to reference sequencing reactions.

### Qualitative and quantitative RT-PCR

For RT-PCR and qRT-PCR analyses of transcript levels, RNA was extracted from strains FA19Str^R^, FA1090, their respective *norM*-negative and *tetR*-negative mutants and FA19Str^R^ _PnorMFA1090_ grown in GCB plus supplements to mid and late logarithmic phases by the TRIzol method as directed by the manufacturer (Thermo Fisher Scientific, Waltham, MA). Genomic DNA (gDNA) was removed by RNAse-free DNAse treatment and gDNA Wipeout (Qiagen, Germantown, MD). The resulting RNA was then reverse transcribed to cDNA using the QuantiTect Reverse Transcriptase kit (Qiagen). Quantitative realtime RT-PCR was performed using the generated cDNA and results were normalized to 16S rRNA expression for each strain. Primers 16Smai_qRTF (5’-CCATCGGTATTCCTCCACATCTCT-3’) and 16Smai_qRTR (5’-CGTAGGGTGCGAGCGTTAATC-3’) were used for the 16S rRNA while primers tetR_qRTR (5’-TTCCACATCAGAGGGCAACA-3’) and tetR_qRTF (5’-GCAACATCAGCACCAAC CAT -3’) were used for the *tetR* gene. Primers N4 and N10 (5’- CATCACGGTATCGACGAAACCGATGCCC-3’) were used for the *norM* gene. Primers murB_qRTF (5’- TAAACACGCCGACGAATTGC-3’) and murB_qRTR (5’-TCTCGCGTA TGCCCTTGTTT-3’) were used for the *murB* gene. All qRT-PCRs were performed in experimental duplicates and biological triplicates. For RT-PCR, random hexamers were used for the reverse transcription while murB_qRTF and tetRSmalR (5’- TCCCCCGGGGGATCGCCCAACAATTCGGCAC-3’), N8 (5’-CCGTTCGGACTGACAGCG-3’) and murB_qRTR were used for PCRs on the cDNA.

### Purification of the TetR protein

Construction of pET15b*tetR* was done by amplifying the *tetR* open reading frame using the primers pETtetR_F (5’- TCGATCCATATGCCCGTGACCCG CATTG-3’) and pETtetR_R (5’-GATCGGATCCTTACGGGTTGCCGTTGCCG -3’). The resulting PCR product along with the pET15b vector were digested with NdeI and BamHI, ligated overnight and transformed into *E. coli* DH5α. The pET15b*tetR* construct was confirmed by sequencing with vector-specific primers T7F (5’-TTAATACGACTCACTATAGG-3’) and T7R (5’-GCTAGTTATTGCTCAGCGG-3’). For protein expression, pET15btetR was transformed into *E. coli* BL21 (DE3) cells. Cultures (5 ml) of BL21 (DE3)-pET15b*tetR* were grown overnight at 30°C and added to 500 ml of LB broth the next morning. The culture was grown at 30°C until mid-log phase and then induced with 0.3 mM IPTG and grown overnight at 30°C. Cells were harvested and resuspended in 20 ml of 10 mM Tris, pH 7.5, 200 mM NaCl, and EDTA-free protease inhibitor was added to bacterial suspension. The cells were lysed by use of a French press cell as described (24), membranes and unbroken cells were removed by centrifugation at 100,000 x *g*, and the supernatant was collected and filtered. TetR-His was purified over a 2-ml nickel-nitrilotriacetic acid (Ni^+2^-NTA) column. After flowing the supernatant over the Ni^+2^-NTA column, the resin was washed successively with buffer containing 20 mM and 50 mM imidazole to remove contaminants and weakly bound proteins, and TetR-His was eluted successively with buffer containing 100 and 200 mM imidazole. The fractions containing TetR-His were concentrated and the imidazole-containing buffer was removed by dialysis into storage buffer (10 mM Tris-HCl, pH 7.5, 200 mM NaCl, and 1 mM EDTA). Dithiothreitol and glycerol were added to a final concentration of 1 mM and 10 % (w/v), respectively.

### Electrophoretic mobility shift assay (EMSA)

DNA probe encompassing the *norM* promoter region was amplified by PCR from FA19 genomic DNA using the upstream primers N11 (5’- CGGTCAGCAGGCGGATTTCTTTCAGG -3’) or N14 (5’-TCTGCCTTCTGTTTTATCCTG -3’). When making radioactive probes, the indicated PCR products were labeled with [γ^32^P]-dATP using T4 polynucleotide kinase (New England Biolabs). The labeled DNA fragments were incubated with 8 μg of TetR-His in 30 μl of reaction buffer at room temperature. For the competition assays, the same non-labelled probe or a non-labelled PCR product using rnpBF1 (5’-CGGGACGGGCAGACAGTCGC-3’) and rnpBR1 (5’- GGACAGGCGGTAAGCCGGGTTC-3’) primers were added in the reaction. Samples were subjected to electrophoresis in a 6% native polyacrylamide gel at 4 °C, followed by autoradiography as described (24).

### Competitive infection of female mice to measure gonococcal fitness

The female mouse model of lower genital tract infection was used to assess whether loss of NorM imposed an in vivo fitness cost or benefit. Mice were inoculated vaginally with an equal number of colony forming units of parent strains FA19Str^R^ and FA1090 with their respective their *norM::kan* transformants and the relative numbers of mutant and wild-type bacteria recovered were compared. The details of the experimental procedures have been described (11, 12, 24). Animal experiments were conducted in the laboratory animal facility at USUHS, which is fully accredited by the Association for the Assessment and Accreditation of Laboratory Animal Care, under a protocol approved by the USUHS Institutional Animal Care and Use Committee.

## ACKNOWLEDGEMENTS

The contents of this article are solely the responsibility of the authors and do not necessarily reflect the official views of the National Institutes of Health, the U.S. Department of Veterans Affairs, or the United States government.

We have no competing interest to declare.

We thank C. del Rio, J. Dillon, R. Nicholas and M. Unemo for providing clinical isolates.

---

**TABLE S1.**
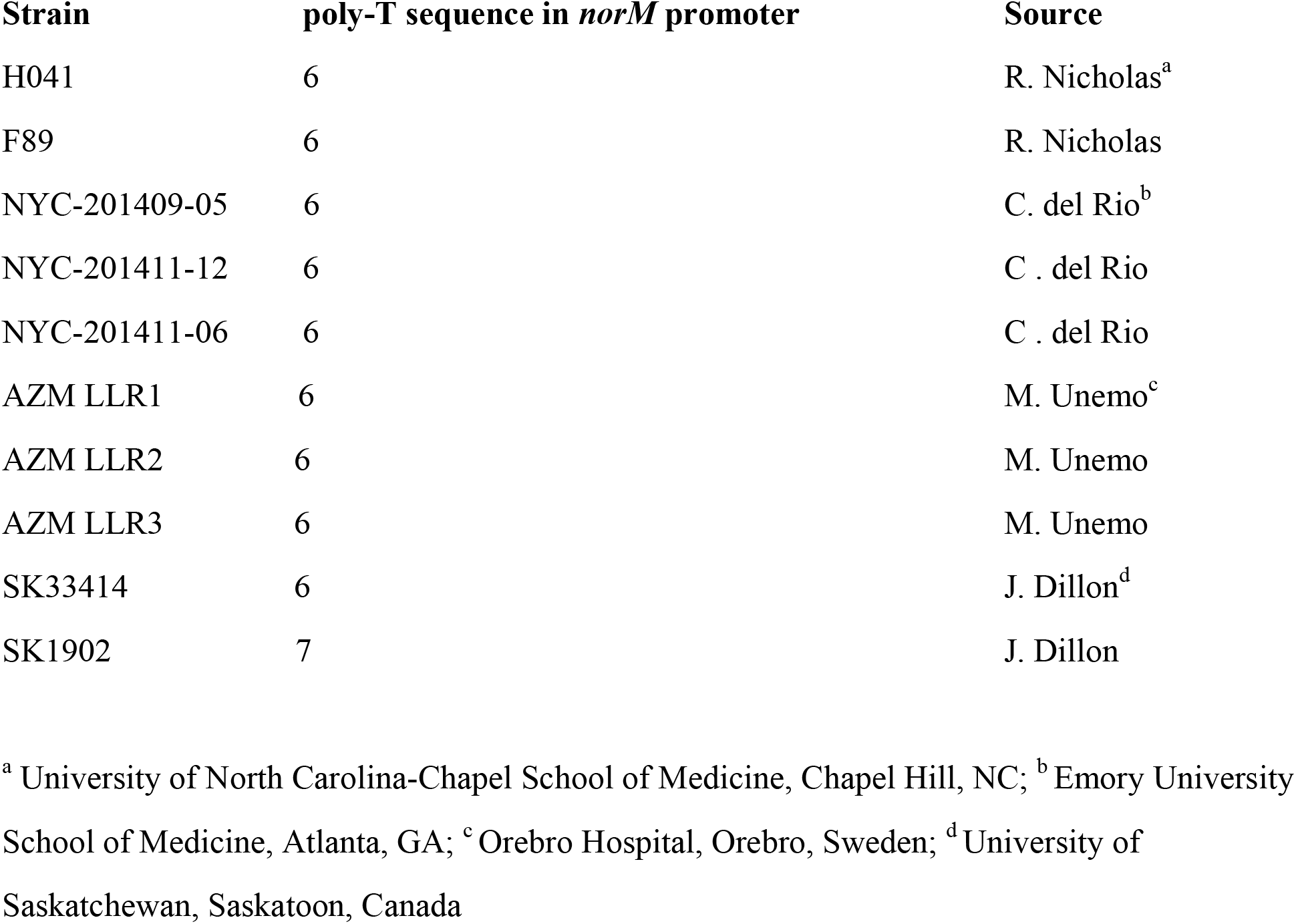

**FIG S1.**
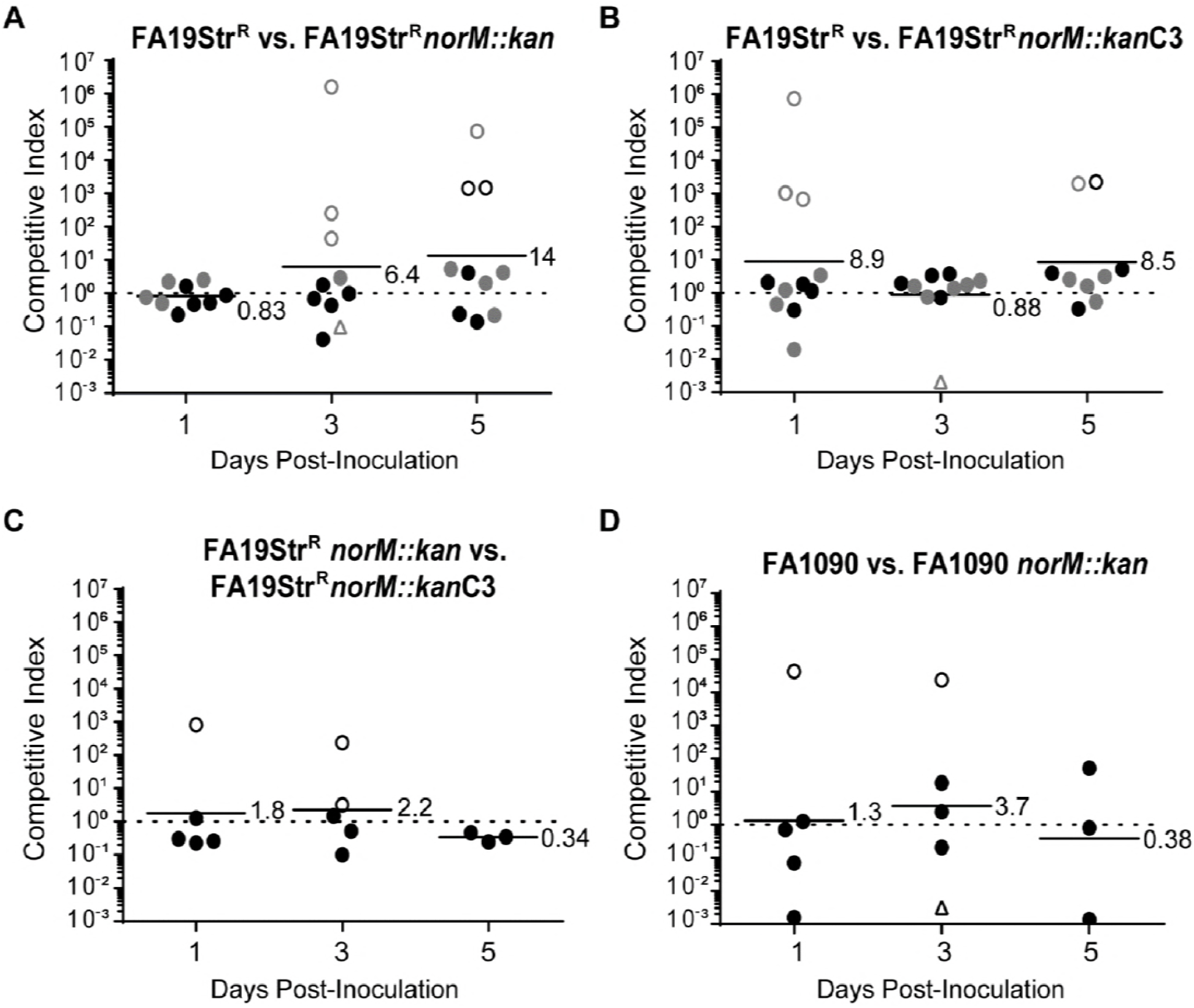
Mutation of *norM* did not alter the *in vivo* fitness of *N. gonorrhoeae* strain FA19 or FA1090 in the female mouse model of gonorrhea infection. Mice were inoculated vaginally with similar numbers of wild-type bacteria and the mutant or complemented mutant strains. Results are expressed as the competitive index (CI) for each mouse on each culture day (CI =1, equal competition; CI <1, mutant strain attenuated; CI >1, mutant strain out-competed the wild-type strain). The geometric mean of the CI values is shown and is represented by the bars. Open circles indicate that only the mutant strain was recovered from the vaginal swabs at the indicated time point. Open triangles indicate that only the wild-type strain was recovered from the vaginal swabs at the indicated time point. A. The FA19Str^R^ *norM::kan* mutant and B. FA19Str^R^ *norM::kan*C3 complemented mutant exhibited similar fitness as wild-type FA19Str^R^ bacteria in vivo. Pairwise analyses of competitive indices from days 1, 3, and 5 of the FA19Str^R^ vs. FA19Str^R^ *norM::kan* and the FA19Str^R^ vs. FA19Str^R^ *norM::kan*C3 competitions did not show a statistical difference by the Mann-Whitney test, indicating that all strains competed similarly *in vivo*. C. No difference in the ability of the FA19Str^R^ *norM::kan* mutant and the FA19Str^R^ *norM::kan*C3 complemented mutant to compete *in vivo* was observed. D. The FA1090 vs. FA1090*norM::kan* competitive experiment also did not show a difference in *in vivo* fitness between the tested strains.

